# Mathematical model of the fast phase of the chlorophyll fluorescence induction curve

**DOI:** 10.1101/2022.08.09.503308

**Authors:** V.M. Lavrentyev

**Affiliations:** V.M. Glushkov Institute of Cybernetics of National Academy of Sciences of Ukraine, 03187, Kyiv, Ukraine

**Keywords:** chlorophyll fluorescence induction, mathematical model, photosynthetic apparatus

## Abstract

In natural conditions, plants are affected by various adverse environmental factors that can disrupt the photosynthetic apparatus, which can reduce the productivity of plants and ultimately reduce their yield. Measurement of the chlorophyll fluorescence induction (CFI) curve is a simple, non-destructive, inexpensive, and fast tool that can be used to analyze photosynthetic reactions and plant conditions. Mathematical modeling of the chlorophyll fluorescence induction curve is important not only for understanding the complex processes of photosynthesis but also can have practical applications in predicting ways to increase plant productivity. Currently, there are a sufficient number of models of varying complexity and detail that describe the processes of photosynthesis, however, no final agreement has been reached. A new model of reactions occurring in the process of the fast phase of the CFI curve, i.e. in the process of electron transport in the electron transport chain (ETC), is presented here. In the ETC model, it is considered as a system of elements in which electrons are sequentially transferred from one element of the system to another, according to the properties of the elements themselves and the connections between them. In addition, the mathematical model is based on the idea of dividing the entire flow of electrons, which moves through the ETC, into a sequence of individual flows. The proposed mathematical model differs in that each stage of electron transfer along the ETC is described separately and sequentially with the help of connection functions. This makes it possible to write the equations for the real OJIP curve and, as a result of their solution, to obtain the parameters of the entire electron transfer process.

---

Research shows that measurements of chlorophyll fluorescence can provide benchmarks to improve agricultural productivity models, increasing the reliability of yield predictions. In natural conditions, plants are exposed to many adverse environmental stressors. They can disrupt the photosynthetic apparatus, causing a decrease in plant productivity and overall yield. Therefore, the measurement of the CFI curve is a simple, non-destructive, inexpensive, and quick tool for analyzing light-dependent photosynthetic reactions and assessing the condition of plants. This makes the method popular among breeders (for example, for crop monitoring), biotechnologists, plant physiologists, farmers, horticulturists, foresters, ecophysiologists, and ecologists [14].

Therefore, mathematical modeling of the CFI curve takes an important place not only for understanding the complex processes of photosynthesis but should take important place and have practical application in predicting possible ways of obtaining higher biomass and productivity of plants [1].

CFI is a manifestation of a very complex biological system, and therefore to describe it correctly and comprehensively is difficult. That is absolutely different from the modeling of technical systems [1].

Despite the complexity of the problem, since the discovery of the fluorescence induction effect of chlorophyll by Kautsky and Hirsch, so many theories have been proposed that more or less describe the processes of photosynthesis in the leaves of higher plants, algae, and cyanobacteria. Naturally, these theories are accompanied by mathematical models. The same applies to such an important process as the process in the electronic transport chain (ETC). The most famous models are described in the literature [2 – 12], and a detailed problems analysis of the photosynthesis modeling is presented in the review [1].

A new model of light-induced photosynthetic reactions occurring during the fast phase of the chlorophyll fluorescence induction curve (OJIP curve) is presented here.

## Separate streams and model structure

The purpose of creating a mathematical model of the fast phase of the chlorophyll fluorescence induction curve (OJIP curve) is to obtain a set of mathematical formulas for describing the processes occurring in the electronic transport chain.

The curve of relative variable fluorescence V(t) = (F(t) – F0) / (FM – F0) was chosen as the basis for modeling, where F(t) is the OJIP curve, F0 is the value of the curve at the starting point, FM is the value curve at the endpoint.

ETC is considered as a system in which electrons are sequentially transferred from one element of the system to others, according to the properties of the elements themselves and the connections between them. Each individual process of electron transfer between two elements of the system is considered an interaction between a source and a receiver and is defined as follows:

- the number of electrons that the source is capable of emitting,
- the number of electrons that the receiver is able to receive, that is, the presence of “vacancies”,
- by the connection function Y(t) = 1 – exp(–k_j_ · t), which abstractly describes the process of electron transfer, where kj determines the rate of electron transfer in a specific j-th source-receiver pair and depends on physicochemical properties and spatial structure of the pair.

The choice of the exponential function is explained by the fact that the exponential nature of various sections of the OJIP curve is noted in many literary sources.

The model does not take into account the back transfers of electrons, which are mentioned in the literature and which are present in other models. If such processes really exist, to simplify the model it is assumed that they do not have a significant effect on the process of direct transfer of electrons by ETC, i.e. they only slow down direct processes.

Fig. 1 shows the structural diagram of the system, which consists of pheophytin Pheo, quinones Q_A_ and Q_B_, a pool of plastoquinons PQ, and cytochrome complex Cyt b6f.

**FIG. 1.**
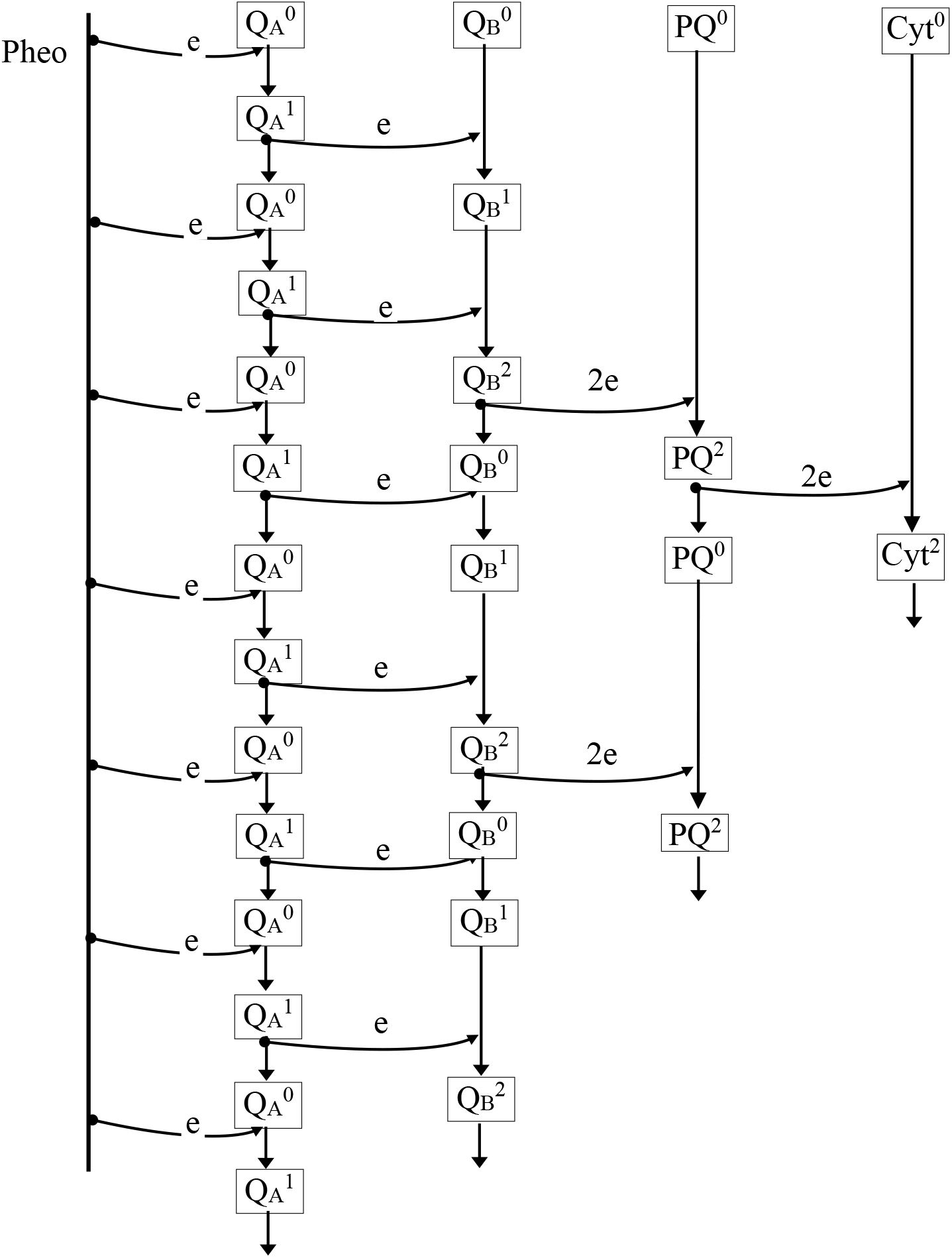
Structural diagram of the model of the electron transfer process by ETL. In the diagram: Pheo – pheophytin, e – electrons, 2e – electron pairs, Q_A_ – one-electron and Q_B_ – two-electron quinone acceptors, PQ – a pool of plastoquinones, Cyt – cytochrome complex.

The mathematical model is based on the idea of dividing the entire flow of electrons, which moves along the ETC, into a sequence of individual flows. According to this, each Q_A_ quinone first accepts the first electron from the first stream from pheophytin. After transferring an electron to the corresponding quinone Q_B_, quinone Q_A_ receives from pheophytin an electron from the second flow, that is, the second electron, then the third, and so on as shown in the diagram.

Processes of acceptance/transfer of electrons by quinones Q_A_ take place sequentially and independently, for example, a single quinone Q_A_ can already receive a second electron, while others have not yet received the first. At the same time, this does not mean that a single Q_A_ quinone cannot receive the second and third electrons until all Q_A_ quinones receive the first.

Since Q_B_ quinones are two-electron acceptors, they successively receive two electrons from Q_A_ quinones and then transfer the pair to the pool PQ. Although the interaction of quinones Q_B_ with the pool PQ is complex, the same exponential function is used in the model.

Next, a pair of electrons is transferred from the pool PQ to the cytochrome complex Cyt b6f. As in the previous case, the exponential function is also used here.

The model also does not take into account the presence of so-called non-regenerating Q_B_ centers, which in one way or another are present in some models.

Electrons gradually fill the “vacancies” of all elements of the system. As shown in the structural diagram of the model, seven streams of electrons are needed to completely fill the ETC with electrons. The complete electron transfer process takes place during the time interval during which the system goes from the initial state in which all Q_A_ quinones are oxidized (“open”) to the final state in which all Q_A_ quinones are reduced (“closed”), which corresponds to the minimum F0 and the maximum FM the value of the fluorescence induction curve.

### Model description

Consider the process of electron transfer by ETC according to the structural model in Fig. 1. It is assumed that the beginning of the OJIP curve is the beginning of the reduction of Q_A_ quinones, that is, after the primary separation of charges with the participation of pheophytin (Р680^*^ Pheo → Р680^+^ Pheo^_^).

For simplification, we will assume that under normal conditions, the number of electrons that pheophytin can provide increases rapidly to the maximum of the relative variable fluorescence function V(t), and is equal to 1. Thus, the function A_1_(t), which determines the number of quinones Q_A_^0^, which receive the first electron from pheophytin depends only on the connection function Y_A1_(t) = 1 – exp(–k_A1_ · t) between pheophytin and quinones Q_A_, i.e.

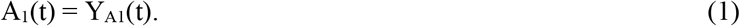

Immediately after the beginning of the reduction of Q_A_ quinones, the process B_1_(t) of the transfer of the first electrons from Q_A_ ^1^ to Q_B_ ^0^ begins. And since all Q_B_ quinones are able to accept electrons, this process will be determined by the number of Q_A_ quinones having the first electron, i.e. A_1_(t) and the connection function Y_B1_(t) = 1 – exp (–k_B1_ · t) between Q_A_ quinones and Q_B_. So

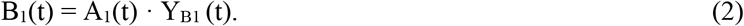

Next, the Q_A_ quinones that transferred the first electron to Q_B_ are reduced a second time. The number of Q_A_ receiving the second electron will be determined by the number of Q_A_ quinones that transferred the first electron to Q_B_ (or by the number of Q_B_ quinones that accepted the first electron, i.e. B_1_(t)), as well as by the connection function Y_A2_(t) = 1 – exp(–k A2 · t) between pheophytin and QA quinones, i.e

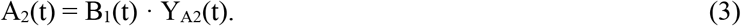

Note that the connection functions Y_A1_(t) and Y_A2_(t) have different coefficients k_A1_ and k_A2_ because the conditions of transfer of the first and second electrons can be different. The same applies to subsequent electron flows.

Immediately after the reduction of Q_A_ quinones, the second electron transfer from Q_A_ ^1^ to Q_B_^1^ begins. This will be determined firstly by the number of Q_A_ that have accepted the second electron, i.e. A_2_(t), secondly by the number of Q_B_ that already have the first electron, i.e. B_1_(t) and thirdly by the connection function Y_B2_(t) = 1 – exp(–k _B2_ · t). According to this, the number of Q_B_ with two electrons will be determined by the function B_2_(t):

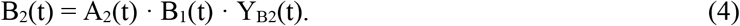

Next, the Q_A_ quinones that have donated a second electron to Q_B_ are reduced a third time. Their number A_3_(t) will be determined by the number of Q_A_ that have transferred two electrons to Q_B_, (or by the number of Q_B_ that have received two electrons from Q_A_, i.e. B_2_(t)), as well as the connection function Y_А3_(t) = 1 – exp(– k_А3_ · t) between pheophytin and Q_A_ quinones. So

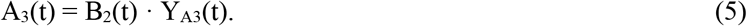

Formulas (1 - 5) describe the transition processes of quinones from the initial state Q_A_^0^ Q_B_^0^ to the state Q_A_^1^ Q_B_^2^. That is, all Q_A_ quinones will be in the Q_A_^1^ state and they will have only the third electron, and all the first and second electrons will be transferred to Q_B_ quinones.

This situation occurs if the transfer of electrons from Q_B_^2^ to the pool PQ is blocked when the plant is treated with methylviologen. This triple reduction of Q_A_ quinones is described as

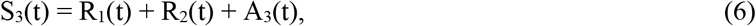

where R_1_(t) = A_1_(t) – B_1_(t) corresponds to the process of acceptance/transfer of the first electrons between quinones Q_A_ and Q_B_, and R_2_(t) = A_2_(t) – B_2_(t) corresponds to the process of acceptance/transfer of the secondary electrons.

These processes are wholly completed when the function A_3_(t) becomes equal to 1, while S_3_(t) corresponds to the relative fluorescence curve when the plant is treated with methylviologen.

If the plant is treated with the drug diuron, the transfer of electrons between quinones Q_A_ and Q_B_ is blocked, and the function A1(t) describes the relative fluorescence curve. If the plant is not subjected to any treatment, the process of transferring electrons in the electronic transport chain continues.

Obviously, as soon as Q_B_ quinones receive a second electron and move to the Q_B_^2^ state, two processes begin. The first is the third reduction of Q_A_ quinones, which have transferred the second electron to Q_B_, according to A_3_(t), and the second is the transfer of electron pairs from Q_B_ quinones to the PQ pool.

Naturally, under normal conditions, the number of transferred pairs of electrons P_1_(t) to the PQ pool will be determined by the number of Q_B_^2^, i.e. B2 (t), as well as the connection function Y_P1_(t) = 1 – exp(–k_P1_ · t) between by quinones Q_B_ and pool PQ.

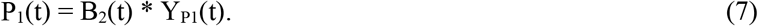

Since at the same time Q_B_^2^ transitions to the Q_B_^0^ state, they begin to accept third electrons from Q_A_, i.e. transition to the Q_B_^1^ state. The number of Q_B_ quinones that receive the third electron will be determined by the number of Q_A_ quinones that have a third electron, i.e. A_3_(t), the number of Q_B_ quinones from which pairs are transferred to the PQ pool, i.e. P_1_(t), and the connection function Y_B3_(t)

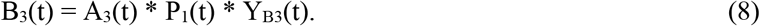

Further, Q_A_ quinones, which transferred the third electron to Q_B_, are restored again - the fourth electron is received from pheophytin. The number of Q_A_ receiving the electron of the fourth stream A_4_(t) will be determined by the number of Q_A_ quinones that have transferred the third electron to Q_B_ (or the number of Q_B_ quinones that have accepted the third electron, i.e. B_3_(t)), as well as the connection function Y_А4_(t) = 1 – exp(–k_А4_ · t)

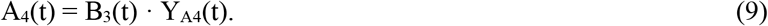

Here we will limit ourselves to the description of the process, given that the structural diagram is simple enough and there is no need to write formulas for other flows, as this should not cause difficulties.

Thus, according to the structural scheme, after the transfer of the seventh electron from pheophytin to Q_A_, the elements of the transport chain enter the states Q_A_^1^, Q_B_^2^, PQ^2^, Cyt^2^. By the way, the seventh flow electron transfer from pheophytin to Q_A_ quinones corresponds to the last rise in the OJIP curve from point I to P.

At this point, the electron transfer process through ETC can be considered complete.

## Results and tasks

This mathematical model differs in that each step of electron transfer along the ETC is described separately and sequentially.

The complete model according to the structure consists of 16 equations (7 equations for Pheo → Q_A_; 6 for Q_A_ → Q_B_; 2 for Q_B_ → PQ; 1 for PQ → Cyt) each of which has an unknown value of k_j_. With the help of these equations, it is possible to describe the entire process of electron transfer through ETC, which corresponds to the curve of relative fluorescence.

It is quite obvious that having the value of the measured real OJIP curve, it is possible to calculate all the values of k_j_ and as a result obtain all the communication functions between the ETC elements.

Fig. 2 shows the curves that correspond to the processes of electron acceptance/transfer between quinones Q_A_ and Q_B_ according to the above formulas 1 – 6. Since these curves are purely illustrative here, the values of k in the formulas are chosen such that the rate of electron acceptance/transfer approximately corresponded to the data given in the literature. Modeling was performed in the MATHCAD system.

**FIG. 2.**
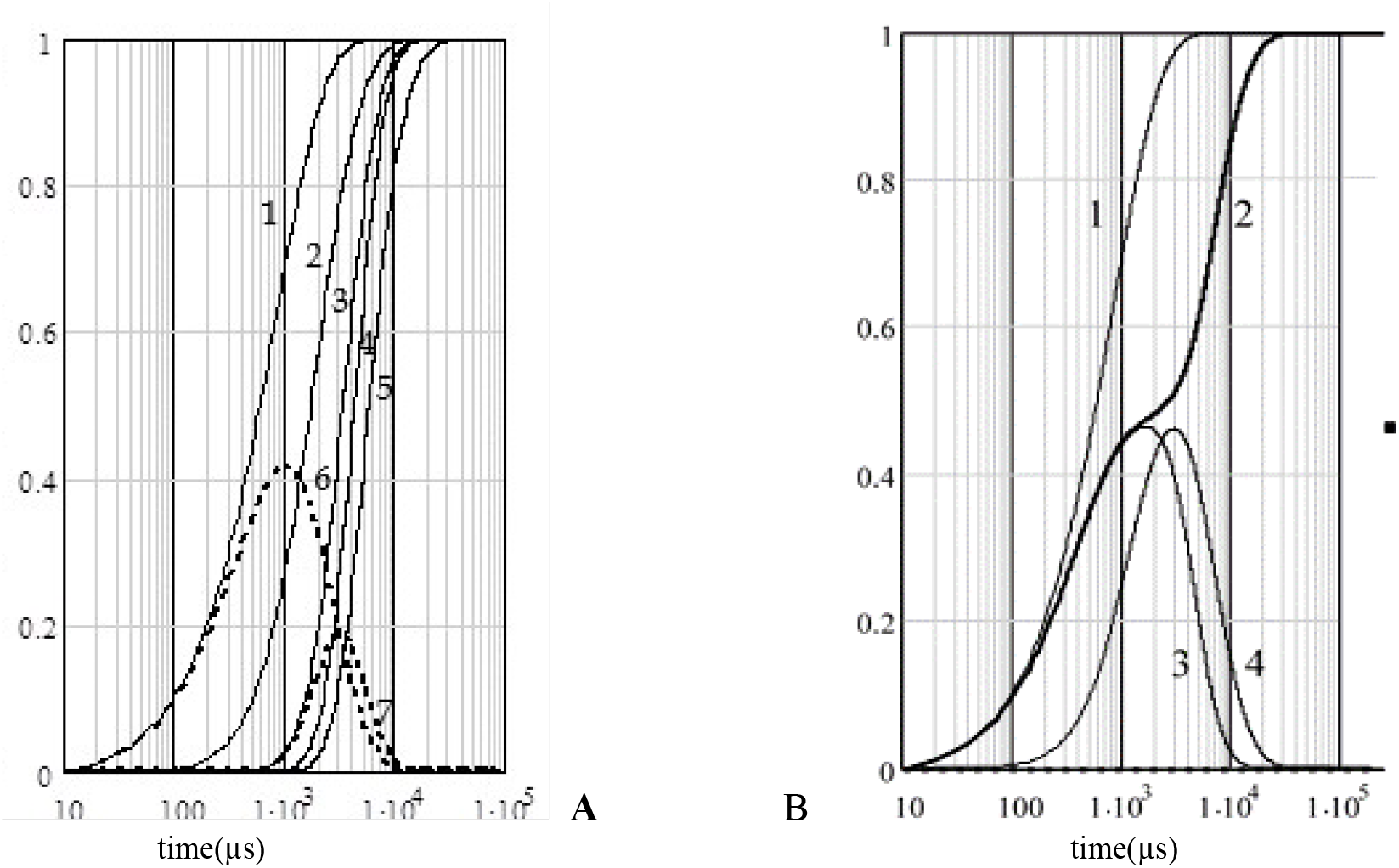
Graphs of functions of electron acceptance/transfer processes by quinones Q_A_ and Q_B_.

Curves 1 – 5 in Fig. 2a are curves A_1_(t), B_1_(t), A_2_(t), B_2_(t) and A_3_(t), respectively. Curve 6 corresponds to the process of receiving/transferring the first electrons between quinones Q_A_ and Q_B_, it is R_1_(t) = A_1_(t) – B_1_(t), and curve 7 is R_2_(t) = A_2_(t) – B_2_(t), which corresponds to the process of acceptance/transfer of second electrons.

Figure 2b shows the following curves: curve 1 is A_1_(t), curve 2 is S_3_(t), curve 3 is the sum of R_1_(t) + R_2_(t), and finally curve 4 is the difference A_1_(t) – S_3_(t). The curve A_1_(t) describes the process of single reduction of Q_A_ quinones in the case when the transfer of electrons from Q_A_ to Q_B_ is blocked, i.e. it corresponds to the fluorescence curve when the plant is treated with diuron. Curve S_3_(t) describes the process of threefold reduction of quinones Q_A_ and each quinone Q_B_ receiving two electrons in the case when the transfer of electrons from Q_B_ to the pool PQ is blocked, i.e. it corresponds to the fluorescence curve when the plant is treated with methylviologen.

Curves 3 and 4 relate to the first rise of the fluorescence curve - OJ. Curve 3 describes the process of receiving/transferring the first and second electrons between quinones Q_A_ and Q_B_. Curve 4 describes the difference between the processes of complete reduction of Q_A_ quinones in “pure” form and reduction of Q_A_ taking into account the transfer of electrons to Q_B_. Considering the biochemistry of the processes of OJ rise, one of these curves can be identified as the one that best corresponds to these processes. And the maximum of the curve will precisely determine the moment of the first inflection of the fluorescence curve, that is, point J.

With the help of this model, it is possible to explain some phenomena of the electron transfer process, which currently do not have a unanimous explanation. For example, when the light flux changes from small values (∼300 μmol m^-2^s^-1^ photons) to (∼3500), the shape of the OJIP curves does not change and is practically proportional to the value of the light flux. When the flow increases to 15,000 μmol m^-2^s^-1^ of photons (Fig. 3А), the shape of the curves changes, and the rate of increase at the beginning of the curve increases, and then a decline appears [11]. This fact can be explained by the fact that a large flow of photons increases the rate of electron acceptance from pheophytin to Q_A_, and the rate of electron transfer from Q_A_ quinones to Q_B_ quinones remains practically unchanged according to the physicochemical properties of the Q_A_ – Q_B_ pair.

**FIG. 3.**
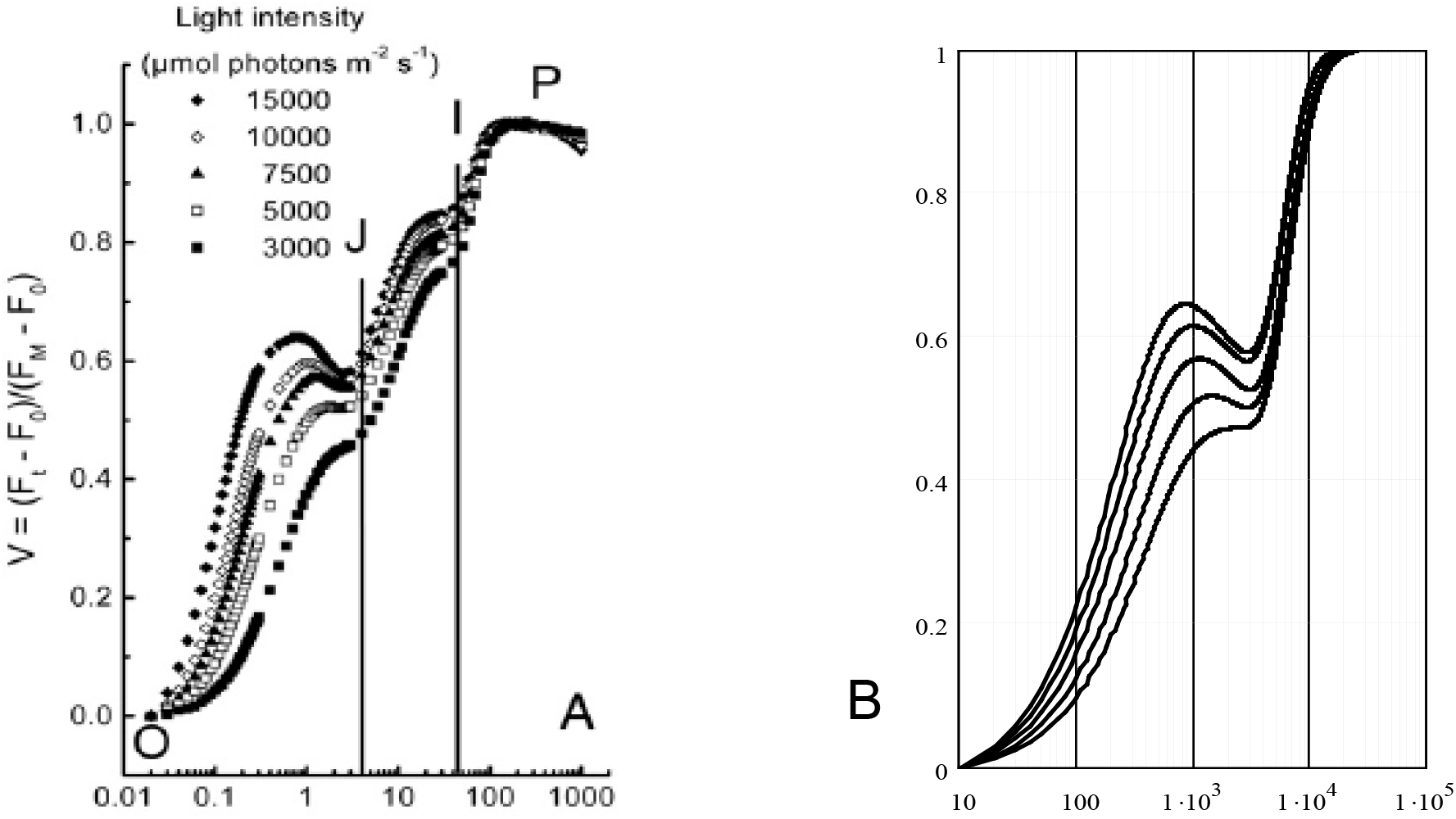
Curves of relative fluorescence at high values of light flux.

Fig. 3b shows the result of modeling a similar change in the shape of the curves at different values of the ratio between k_B_ and k_A_ at constant k_B_.

The mathematical model was tested on several OJIP curves. The closeness of the simulated curves to the real ones was 98-95% (test results are not shown due to the small number of test curves).

The proposed model can be very useful for analyzing the impact of stress factors on the condition of plants. The review [14] gives examples of the influence of temperature and water factors, salt, nitrates and heavy metals on plants. Conclusions about the presence of stress are made based on the analysis of the parameters of the OJIP curve, that is, the values at the characteristic points O, J, I, P and the distance between them. Without resorting to the positive and negative sides of such an analysis, let us note only that the values at points J and I are the result of several simultaneous processes and changes in one place can be compensated for in another, which will complicate the analysis.

Since in the proposed model the processes in ETC are separated from each other, the influence of the stress factor can be detected at an early stage if there is an analysis of a sufficient database of measurements of real OJIP curves. This can be very effective in digital farming and greenhouses where constant monitoring and feedback is performed.

## Conclusions

The proposed mathematical model differs in that each stage of electron transfer along the ETL is described by the connection functions between ETL components and the idea of dividing the entire electron flow into separate flows.

The model makes it possible to write a system of equations for the OJIP curve and obtain the entire curve parameters and individual stages resulting from the equation’s solution.

The structure of the model is very flexible and allows you easily replace any connection function. Such a need may arise when determining another connection function due to research. In addition, it is possible to conduct purely theoretical experiments - to determine how the shape of the OJIP curve changes depending on the connection functions and their parameters.

The proposed mathematical model can provide many answers to the photosynthesis questions, but it is necessary to carry out additional research. First, to optimize the solving algorithm of the equations system and determine the minimum but a sufficient number of curve points and their location on the time axis.

It is also necessary to analyze the model for the fast algorithm development for determining stress states based on measurement data.

